## Supplemental Data for "A β-Galactosidase acting on unique galactosides: the structure and function of a β-1,2-galactosidase from *Bacteroides xylanisolvens*, an intestinal bacterium"

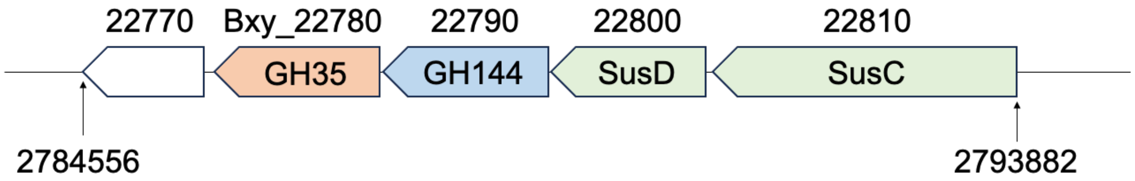

**Figure S1. The gene cluster around the gene encoding GH144 enzyme in *B. xylanisolvens* genome.** Genes and their directions are shown by arrows. Annotation of functions of the genes are represented by color in the arrows: GH35, light orange; GH144, light blue; SusCD, light green. The gene loci (KEGG database, https://www.genome.jp/kegg/) are shown above the arrows, and "Bxy_" are omitted except Bxy_22780. The numbers below the genes are nucleotide numbers from 5´ of the genomic DNA.

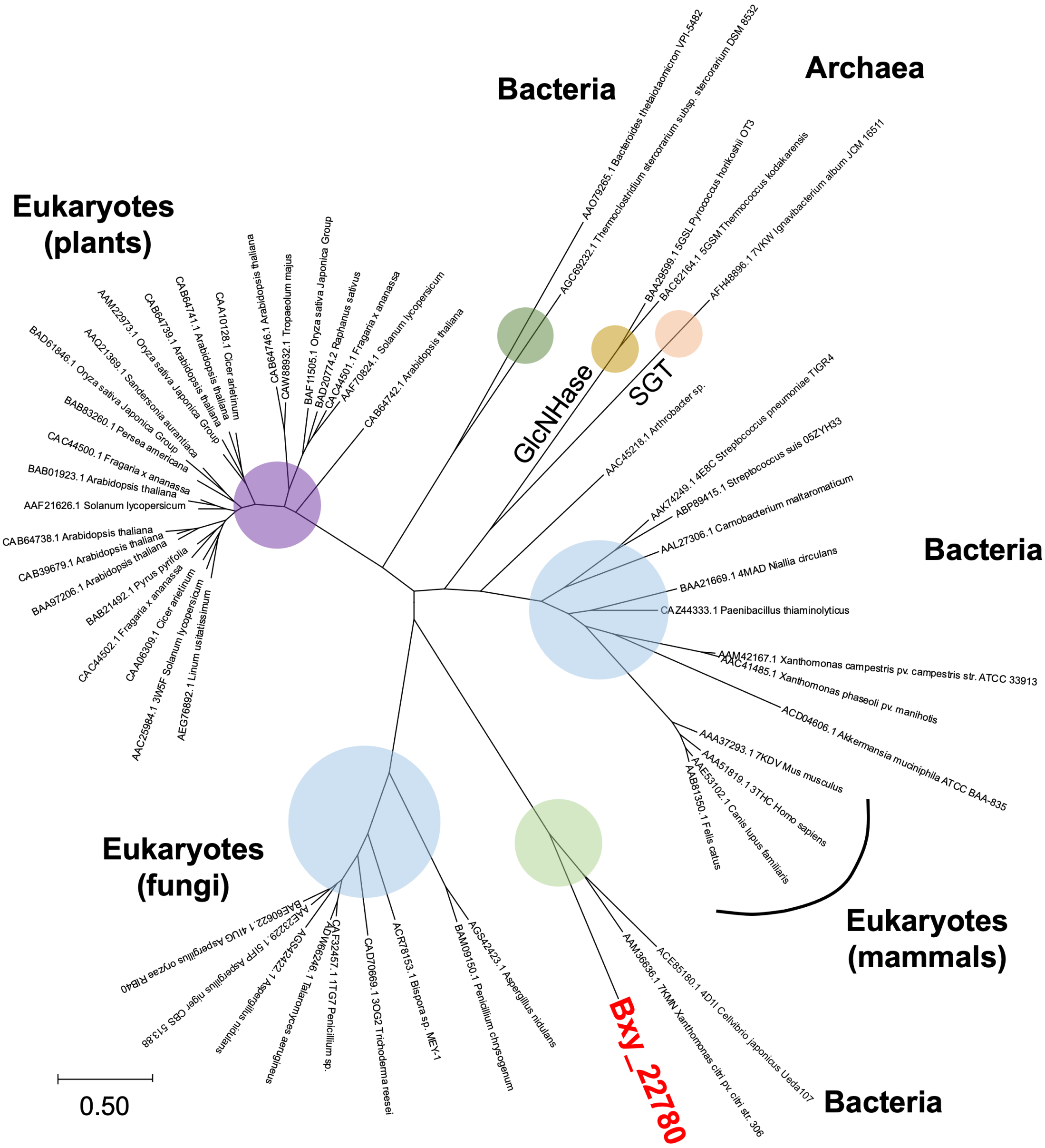

**Figure S2. Phylogenetic tree of GH35 enzymes**

Bxy_22780 is shown in red. The other proteins are indicated with GenBank accession No., organism names, and PDB IDs (if available). GlcNHase and SGT are abbreviations of β-glucosaminidase and β-1,2-glucosyltransferase, respectively. All except GlcNHases and SGT (enzymes without enzyme names) are β-galactosidases. A scale bar is shown at bottom left of the figure. Phylogenetic groups are visualized by colored circles.

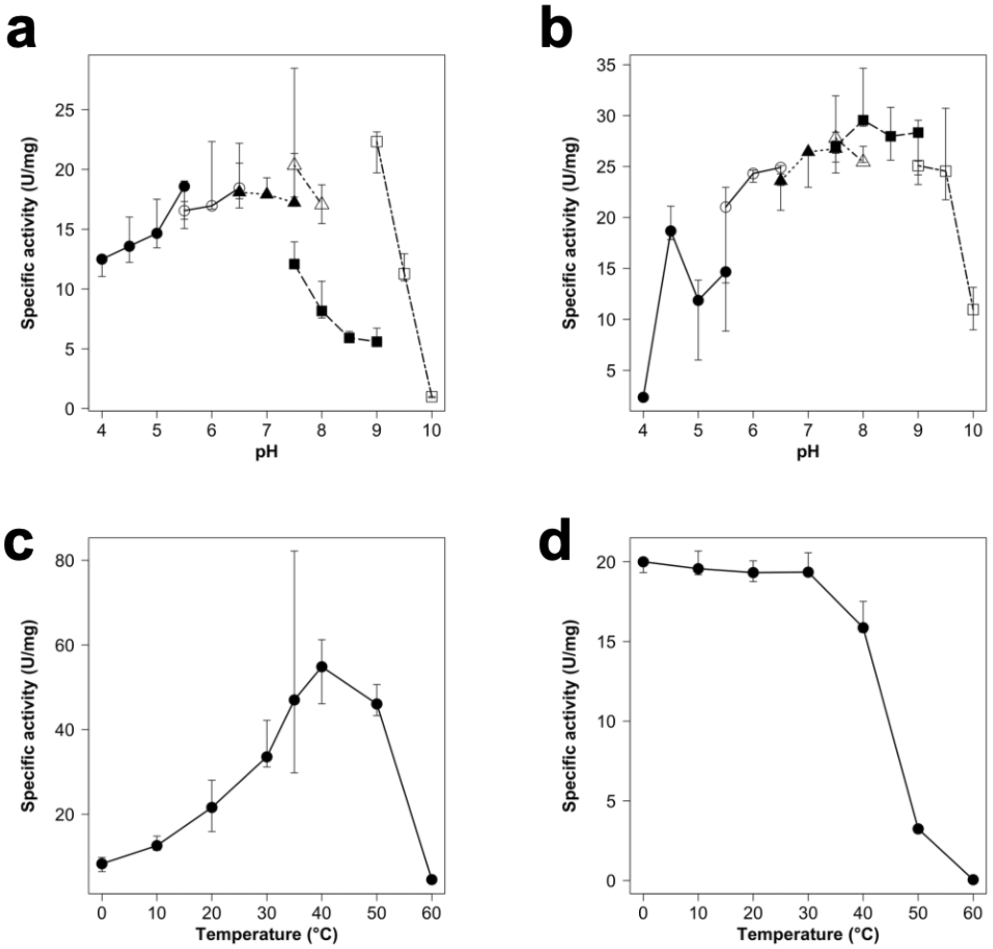

**Figure S3. pH and temperature profiles of Bxy_22780**

**a–b,** Optimum pH (**a**) and pH stability (**b**). Buffers used are sodium acetate (pH 4.0–5.5, closed circles), MES (pH 5.5–6.5, open circles), MOPS (pH 6.5–7.5, closed triangles), HEPES (pH 7.5–8.0, open triangles), Tris-HCl (pH 7.5–9.0, closed squares) and glycine (pH 9.0–10.0, open squares). **c–d**, Optimum temperature (**c**) and temperature stability (**d**). Medians in triplicate experiments are shown as plots and the other data are shown as error bars.

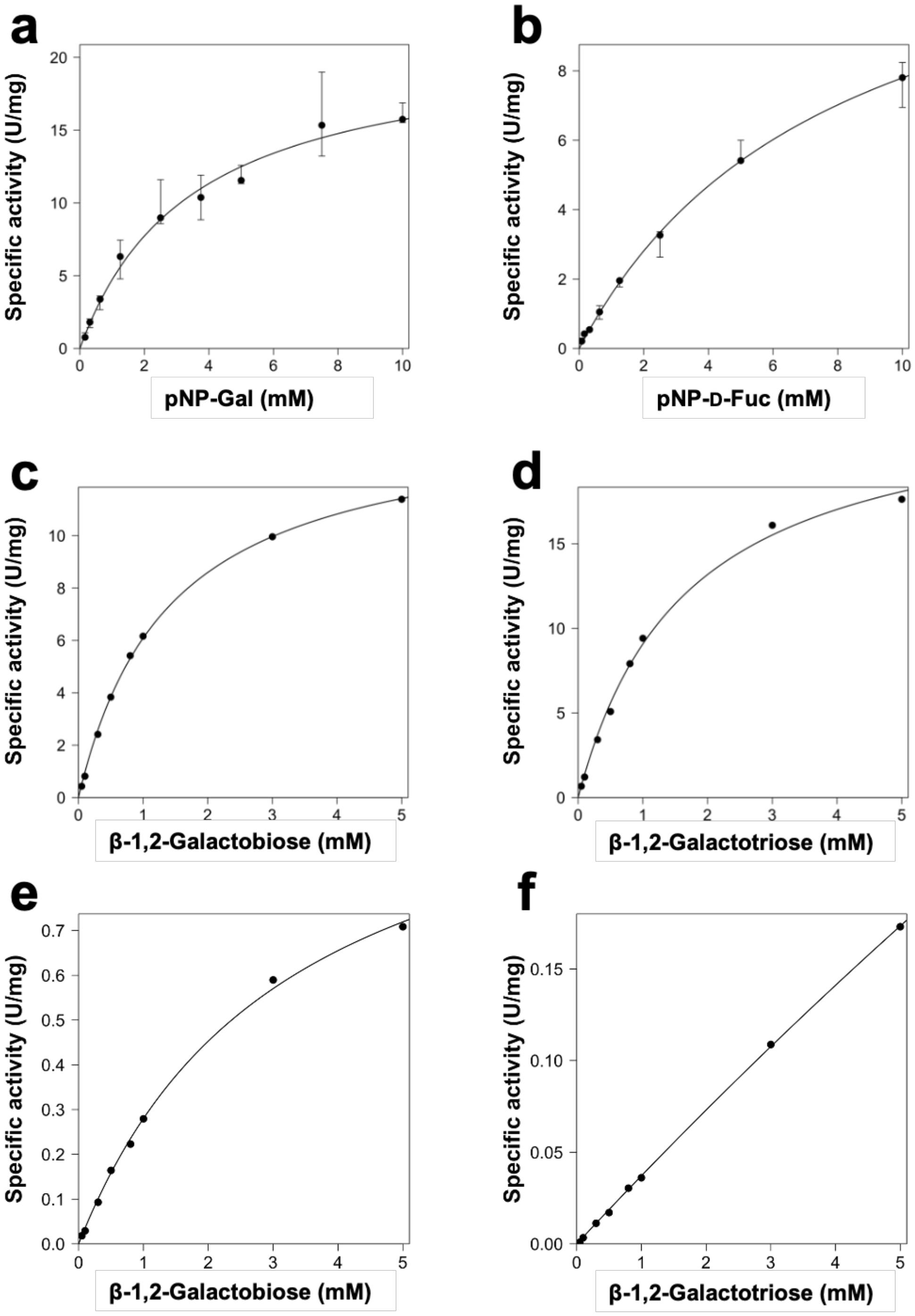

**Figure S4. Kinetic analysis of Bxy_22780**

Kinetic analysis for the wild-type (**a**–**d**) and W288A mutant (**e**–**f**). Experiments were performed triplicate (**a**–**b**) for pNP-sugars and once (**c**–**f**) for β-1,2-galactooligosaccharides. Maximum and minimum data in the triplicate are used as error bars (**a**–**b**). Kinetic parameters are shown in Table 1.

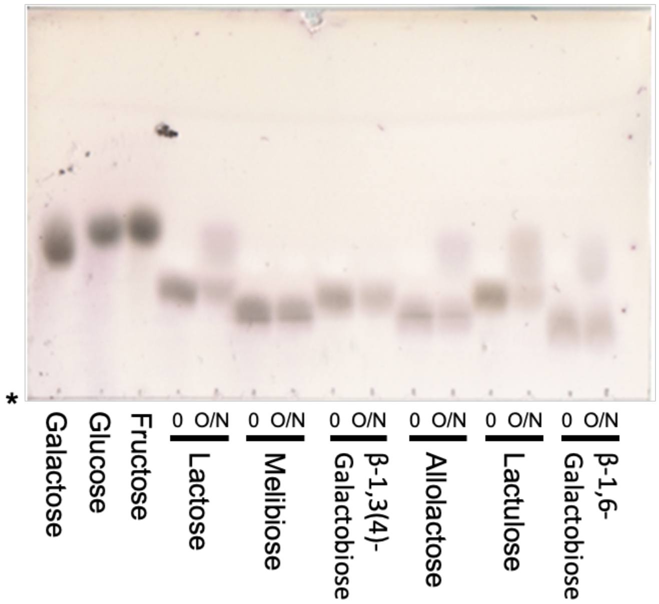

**Figure S5. Hydrolytic activity of Bxy_22780 toward galactosides**

Each reaction was performed in the presence of 5 mM substrate and 0.1 mg/ml Bxy_22780 (wild-type). An asterisk represents an origin. Lanes Fructose, glucose and galactose are markers (1 μl of 10 mM solution). Reaction time are shown as “0” and “O/N” for 0 h and overnight, respectively.

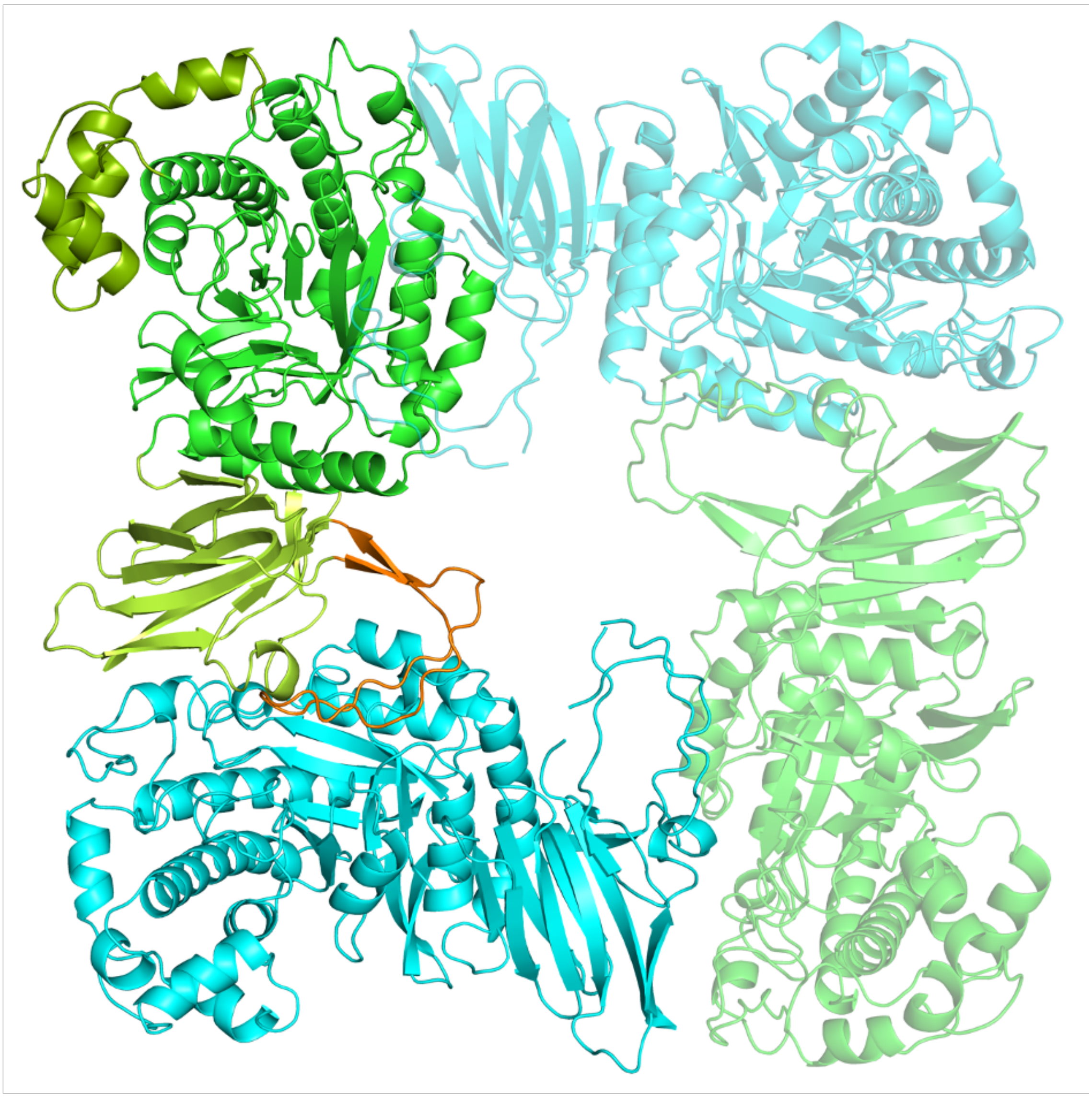

**Figure S6. Overall structure of Bxy_22780**

PDB ID of the overall structure of E350G mutant is 8Z43. The two subunits in an asymmetric unit are fully colored while the other subunits in a symmetry mate are shown transparently. The two subunits in the asymmetric unit are colored in green and cyan basically. In the green colored subunit, a catalytic domain composed of TIM-barrel domain (residues 1–198 and 253–404) are left in green while an inserted region in the catalytic domain (residues 199–252), DUF5597 domain (residues 405–506 and 539–550), and an inserted region in DUF5597 domain are colored in dark green, light green and orange, respectively.

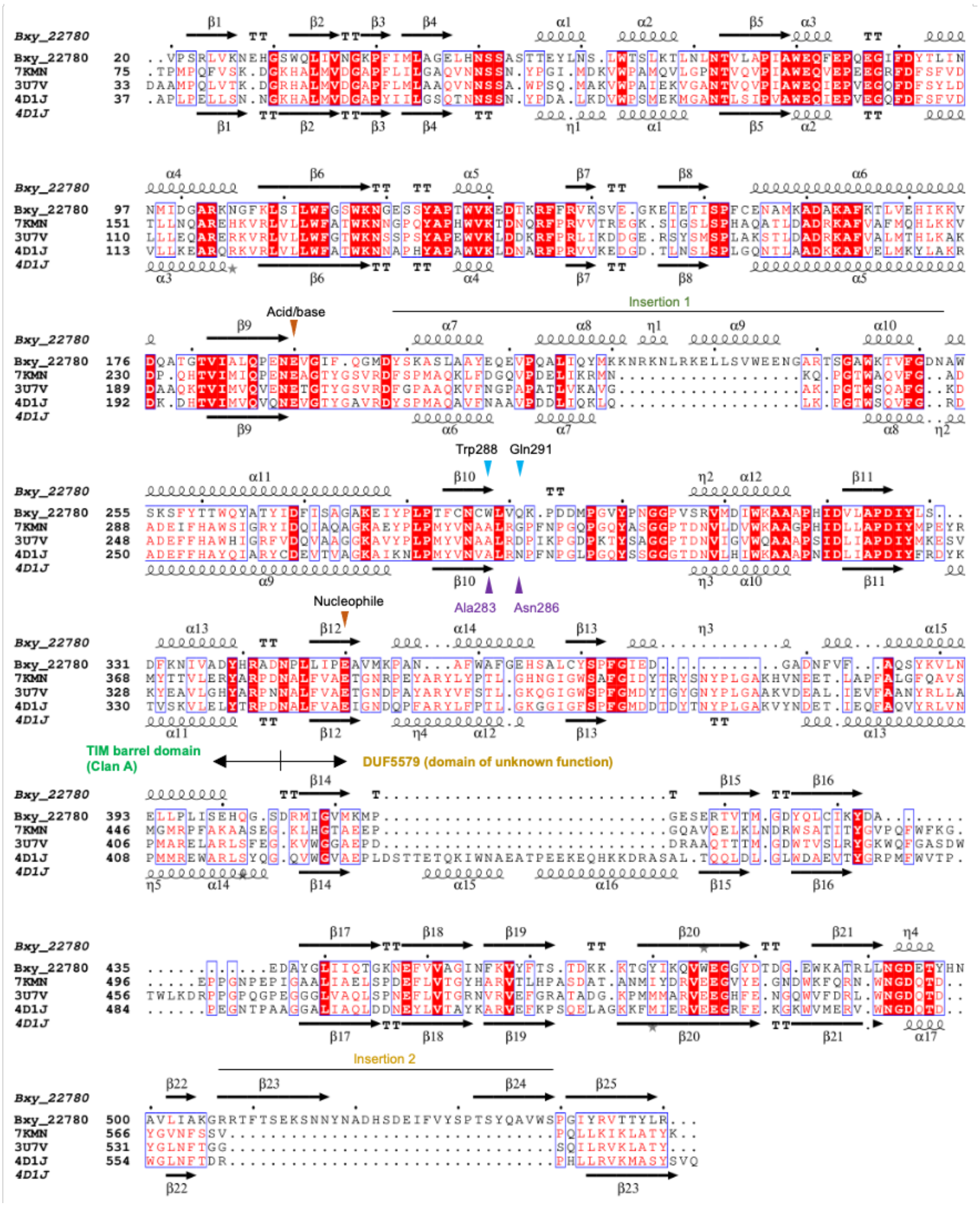

**Figure S7. Structure-based multiple sequence alignment**

Multiple sequence alignment was performed using PDBeFold (https://www.ebi.ac.uk/msd-srv/ssm/) [1]. The alignment was visualized by ESPript ver3.0 (https://espript.ibcp.fr/ESPript/cgi-bin/ESPript.cgi) [2]. The constitution of the domains and insertion regions shown in Fig. S6 are added manually in the alignment. Catalytic residues (Glu190 and Glu350) and substrate recognition residues (Trp288 and Gln291) in Bxy_22780 are indicated with dark orange and cyan reversed triangles, respectively. The

Ala283 and Asn286 in CjBgl35A corresponding to Trp288 and Gln291 are indicated with purple triangles. The aligned amino acid sequences other than Bxy_22780 (GenBank accession NO., CBK67349.1) are shown as PDB IDs: 7KMN (*Xanthomonas citri* pv. *citri* str. 306, AAM36636.1), 3U7V (*Caulobacter vibrioides* CB15, AAK22773.1) and 4D1J (*Cellvibrio japonicus* Ueda107, ACE85180.1).

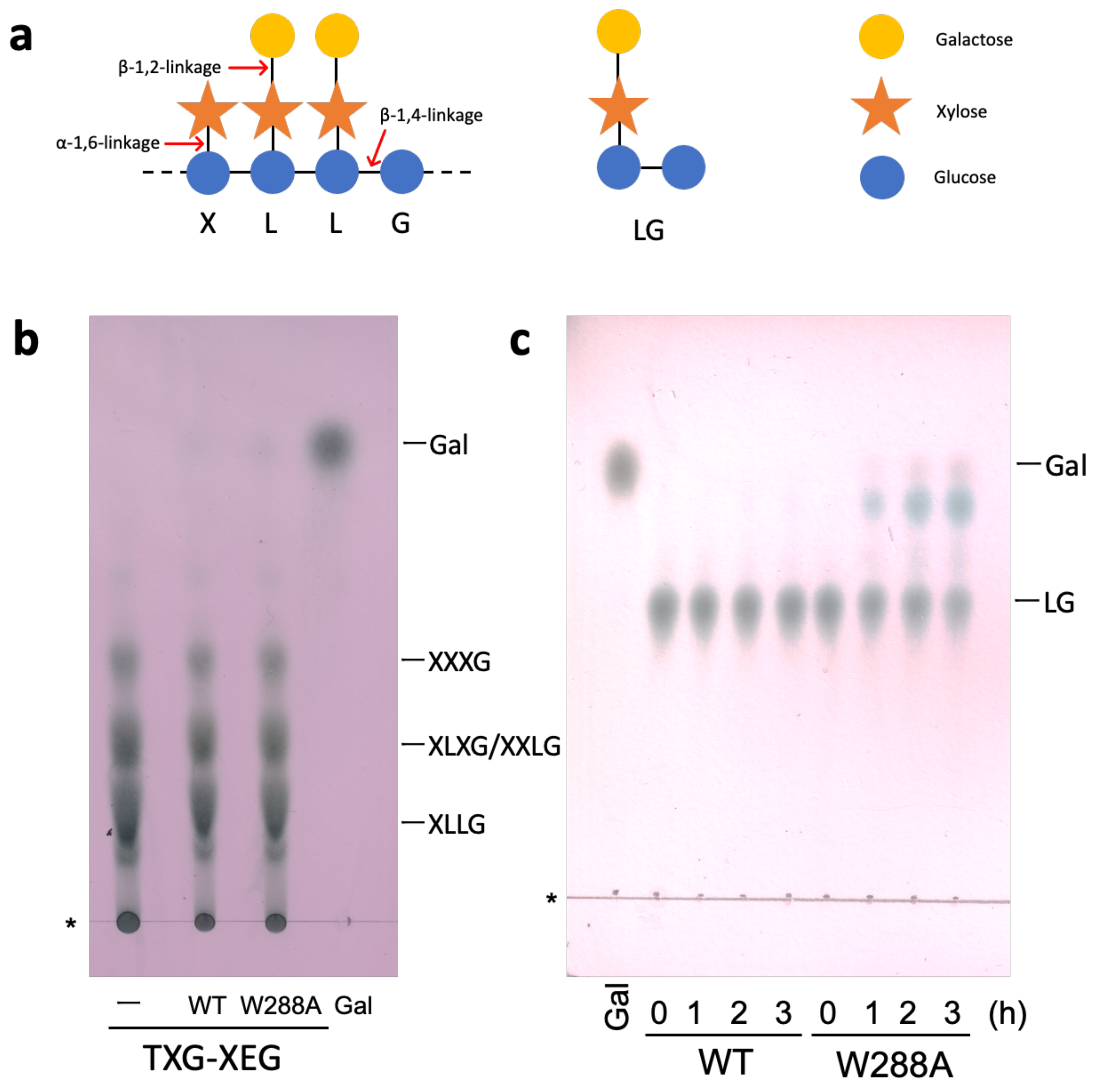

**Figure S8. Activity toward xyloglucan associated substrates**

**a**, Schematic representation of tamarind-xyloglucan. Representation of side chain patterns in xyloglucan is used in **b**, **c**: G, no side chain; X, with xylose as a side chain; L, galactosyl xylose as a side chain. Symbols of monosaccharides illustrated based on Symbol Nomenclature for Glycans (https://www.ncbi.nlm.nih.gov/glycans/snfg.html). **b** and **c**, Action pattern analysis toward TXG-XEG (**b**) and LG (**c**) by TLC. Lane Gal, 10 mM Gal (1 μl) was spotted. Lane −, TXG-XEG without enzyme. Asterisks represent the origin on the TLC plates.

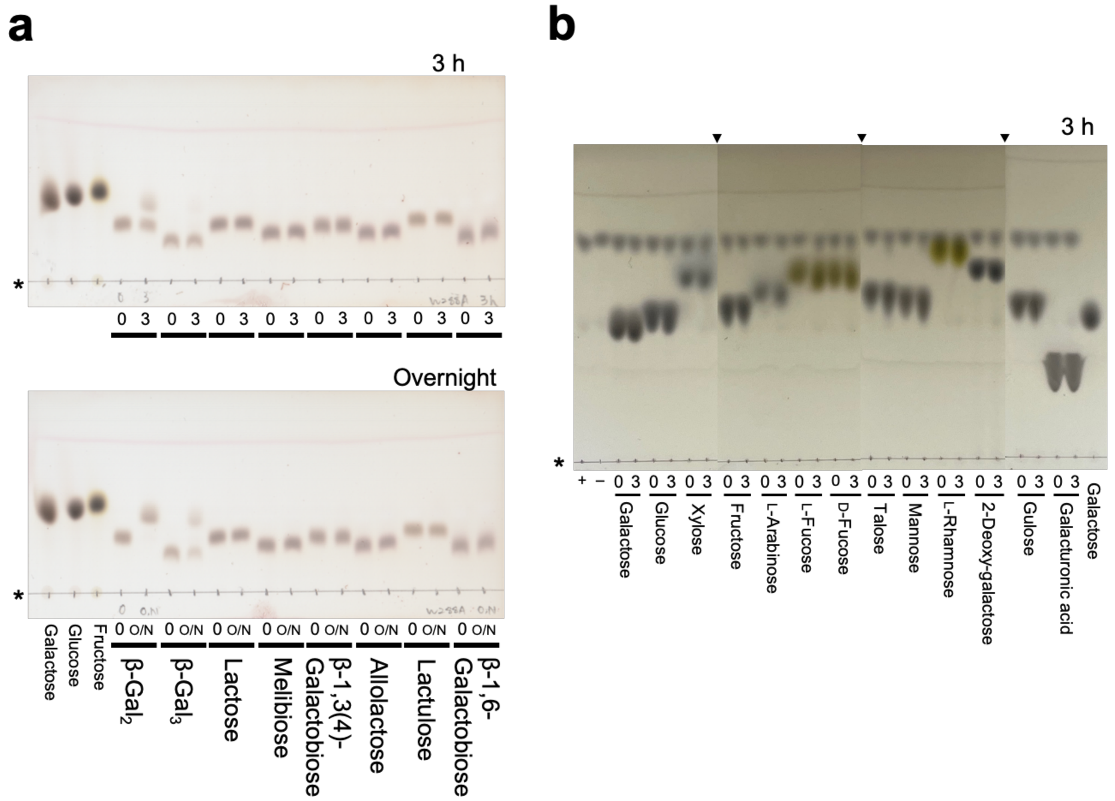

**Figure S9. Substrate specificity of W288A mutant**

**a**, Hydrolytic activity of W288A mutant. **b**, Glycosynthase activity of W288A/E350G mutant. Lanes without bars are markers (1 µl of 10 mM solution). Lanes indicated with + and – represent αGalF was incubated with and without W288A/E350G mutant, respectively. Reaction times are shown above the bars and the plates. Reverse triangles above the plates represent the boundaries of TLC plates.

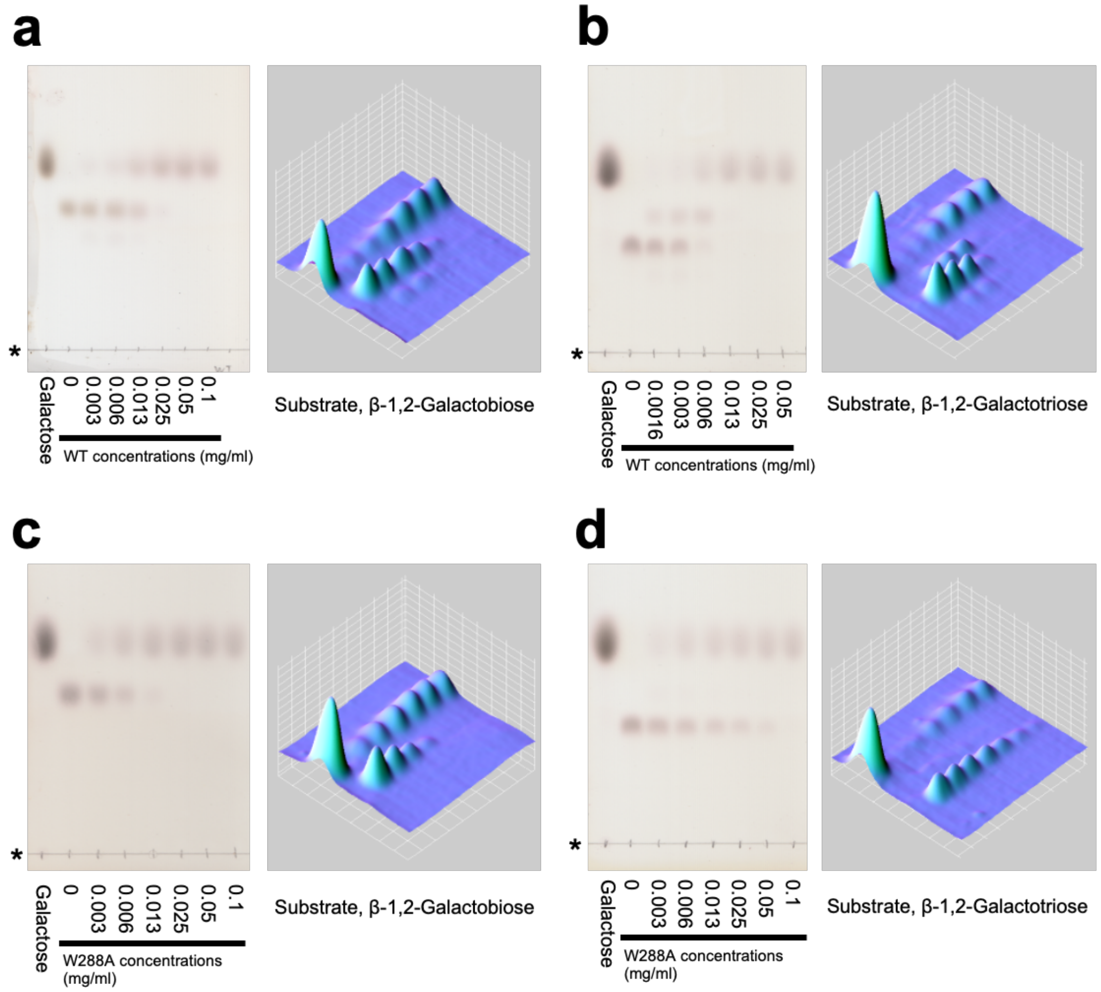

**Figure S10. Transglycosylation activity toward β-1,2-galactooligosaccharides**

(Left) Action patterns shown by TLC plates. Lane Galactose, 1 µl of 10 mM galactose was spotted. Asterisks represent the origins. (Right) Graphic visualization of the TLC plates by ImageJ [3].

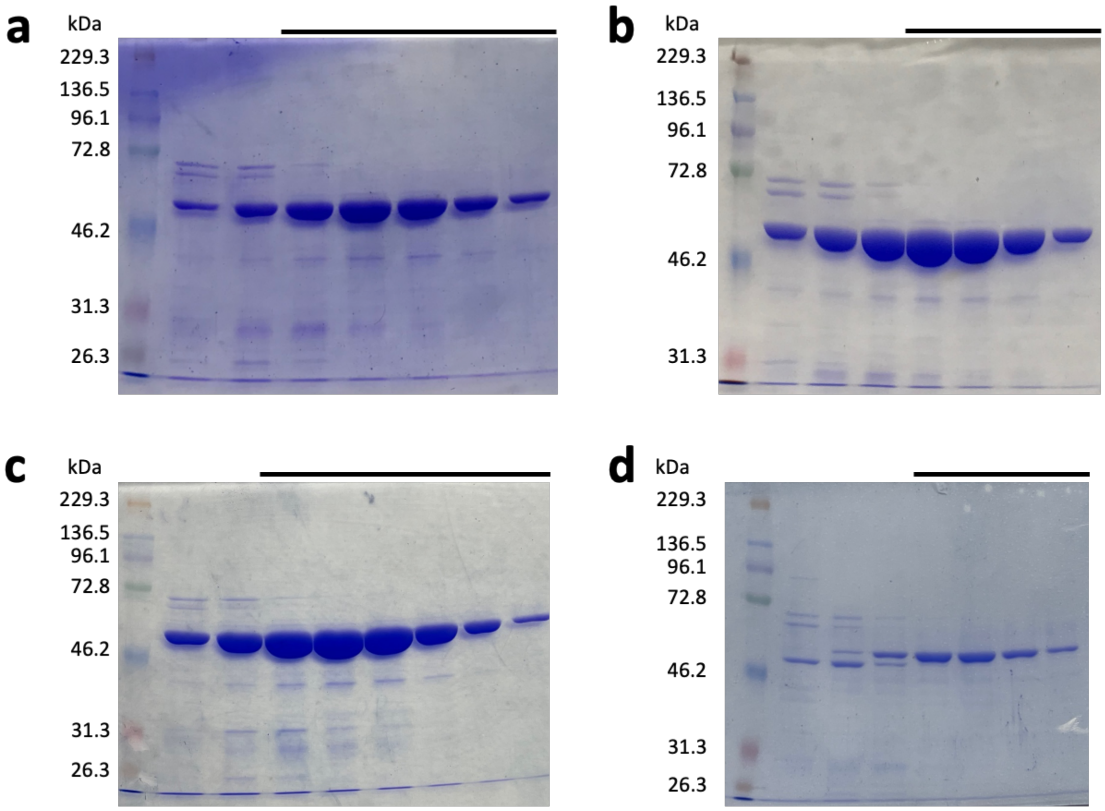

**Figure S11. SDS-PAGE of Bxy_22780**

Fractions obtained using Nickel affinity column for purification of the wild-type (**a**), E350G mutant (**b**), W288A mutant (**c**) and W288A/E350G mutant (**d**). Bars above the gels represent the fractions collected for further use. Left lanes are marker proteins.

**Supplementary Table 1. Chemical shifts in $^{13}$C NMR and $^{1}$H-NMR spectra of β-1,2-$Gal_2$**

| Sugar ring[a] | Position | α | | | | β | | | |
|---|---|---|---|---|---|---|---|---|---|
| | | $^{13}$C NMR (δ ppm) | $^{1}$H NMR (δ ppm) | | *J* (Hz) | $^{13}$C NMR (δ ppm) | $^{1}$H NMR (δ ppm) | | *J* (Hz) |
| I | 1 | 91.8 | 5.47 | d[b] | *J* = 4.0 | 94.8 | 4.66 | d | *J* = 8.0 |
| | 2 | 77.9 | 3.88–3.90 | m | | 79.7 | 3.72 | m | |
| | 3 | 67.8 | 4.01 | m | | 72.5 | 3.82–3.83 | m | |
| | 4 | 69.0 | 4.02 | m | | 68.5 | 3.93–3.99 | m | |
| | 5 | 69.9 | 4.09 | m | | 75.0 | 3.71 | m | |
| | 6 | 60.8 | 3.74 | m | | 60.6 | | | |
| II | 1 | 104.3 | 4.55 | d | *J* = 7.6 | 103.1 | 4.67 | d | *J* = 8.0 |
| | 2 | 70.7 | 3.61 | m | | 71.1 | 3.56 | m | |
| | 3 | 72.3 | 3.64 | m | | 72.5 | 3.61–3.64 | m | |
| | 4 | 68.3 | 3.90–3.91 | m | | 68.4 | | | |
| | 5 | 74.7 | 3.67–3.69 | m | | 74.9 | 3.75–3.77 | m | |
| | 6 | 60.6 | 3.77–3.78 | m | | 60.9 | 3.65–3.66 | m | |

[a] I and II denote the first and second galactoside units from reducing end, respectively.

[b] The signals are described as d = doublet; m = multiplet.

**Supplementary Table 2. Data collection and refinement statistics**

| Data set | Ligand-free (E350G) | WT-Gal | WT-MeβGal |
|---|---|---|---|
| **Data collection** | | | |
| Beamline | KEK BL-5A | KEK BL-5A | KEK BL-5A |
| Space group | *I* 2 | *I*4 | *C*2 |
| Unit cell parameters (Å, º) | *a* = 144.70 Å<br>*b* = 51.06 Å<br>*c* = 180.08 Å<br>β = 90.41 º | *a* = 163.83 Å<br>*b* = 163.83 Å<br>*c* = 50.86 Å | *a* = 231.76 Å<br>*b* = 50.84 Å<br>*c* = 229.45 Å<br>β = 96.10 º |
| Resolution (Å)[a] | 48.198–1.91 (1.94–1.91) | 41.78–1.86 (1.90 –1.86) | 45.91–1.94 (1.97–1.94) |
| Total reflections[a] | 680380 (34802) | 377013 (23608) | 601647 (29660) |
| Unique reflections[a] | 102654 (5058) | 57035 (3503) | 194503 (9674) |
| Completeness (%)[a] | 99.9 (100) | 100 (99.9) | 98.2 (99.5) |
| Multiplicity[a] | 6.6 (6.9) | 6.6 (6.7) | 3.1 (3.1) |
| Mean $I/\sigma(I)$[a] | 15.8 (2.2) | 10.3 (2.2) | 8.1 (2.0) |
| $R_{merge}$ (%)[a] | 8.5 (78.8) | 9.6 (79.2) | 7.1 (48.4) |
| $R_{pim}$ (%)[a] | 5.4 (49.2) | 6.1 (49.5) | 6.9 (46.7) |
| $CC_{1/2}$[a] | (0.754) | (0.711) | (0.768) |
| **Refinement** | | | |
| Resolution (Å) | 48.152–1.91 | 41.779–1.86 | 45.629–1.94 |
| No. of reflections | 97427 | 54141 | 184670 |
| No. of atoms | 18328 | 8649 | 36003 |
| No. of water molecules | 528 | 223 | 1329 |
| $R_{work}/R_{free}$ (%) | 17.4/20.2 | 19.3/22.7 | 21.1/25.1 |
| No. of asymmetric units | 2 | 1 | 4 |
| RMSD from ideal values | | | |
| Bond lengths (Å) | 0.0081 | 0.0084 | 0.0070 |
| Bond angles (°) | 1.6821 | 1.7231 | 1.5352 |
| Average *B*-factors (Å$^2$) | | | |
| Protein (chain A/B/C/D) | 29.2/29.4 | 34.0 | 26.4/25.9/27.4/26.5 |
| Ligand | | | |
| Gal or MeβGal (chain A/B/C/D) | - | 35.1 | 29.1/31.1/31.4/29.5 |
| Solvent | 31.3 | 35.2 | 30.0 |
| Ramachandran plot (%) | | | |
| Favored | 98.0 | 98.0 | 98.0 |
| Allowed | 2.0 | 2.0 | 2.0 |
| Outlier | 0 | 0.0 | 0 |
| **PDB entry** | 8Z43 | 8Z47 | 8Z48 |

[a] Values in parentheses represent the highest resolution shell.

**Supplementary Table 3. Primers used in this study**

| | Sequence (5´ to 3´) | |
|---|---|---|
| Subcloning of the whole gene in databases | | |
| Fw | TTTCATATGGGAACTTCAATGATGAATG | NdeI |
| Rv | TATGAATAAGCTGCTCGAGCCGTAAATAGGTTGTC | XhoI |
| Deletion of the N-terminal signal peptide | | |
| Fw | ACATATGCGCCCCCCAATGTATACT | |
| Rv | GGGGGGCGCATATGTATATCTCCTTC | |
| Mutation of E350G | | |
| Fw | ATTCCCGGTGCAGTGATGAAACCGGCC | |
| Rv | CACTGCACCGGGAATCAATAACGGATT | |
| Mutation of W288A | | |
| Fw | AACTGTGCACTGGTGCAAAAGCCTGAT | |
| Rv | CACCAGTGCACAGTTGCAGAAGGTAGG | |
| For assay | | |
| TM1867 (L-lactate dehydrogenase) | | |
| Fw | GTGATTCATATGAAAATAGGTATCG | NdeI |
| Rv | CTCTACCTCGAGTTAACCGCTGGTG | XhoI |
| TM1190 (galactokinase) | | |
| Fw | TTTAAGAAGGAGATATACATATGAAAGTGAAGGCACCAGGAAG | |
| Rv | GTGGTGGTGGTGGTGGTGCTCGAGGATTTTTGAACACCGTCTGAAC | |

Restriction sites are indicated with underlines and restriction enzymes acting on the underlined sequences are shown beside the sequences. Fw, forward primer; Rv, reverse primer.

### Table of Contents

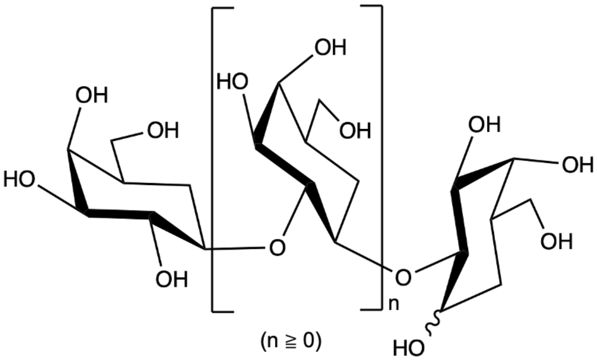

β-1,2-Galactooligosaccharide

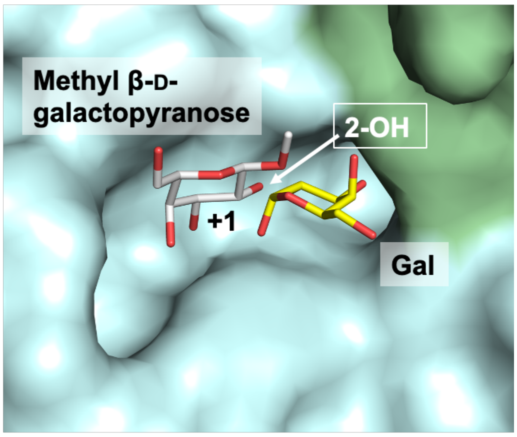

**β-1,2-Galactosidase**

The enzyme with a new catalytic reaction

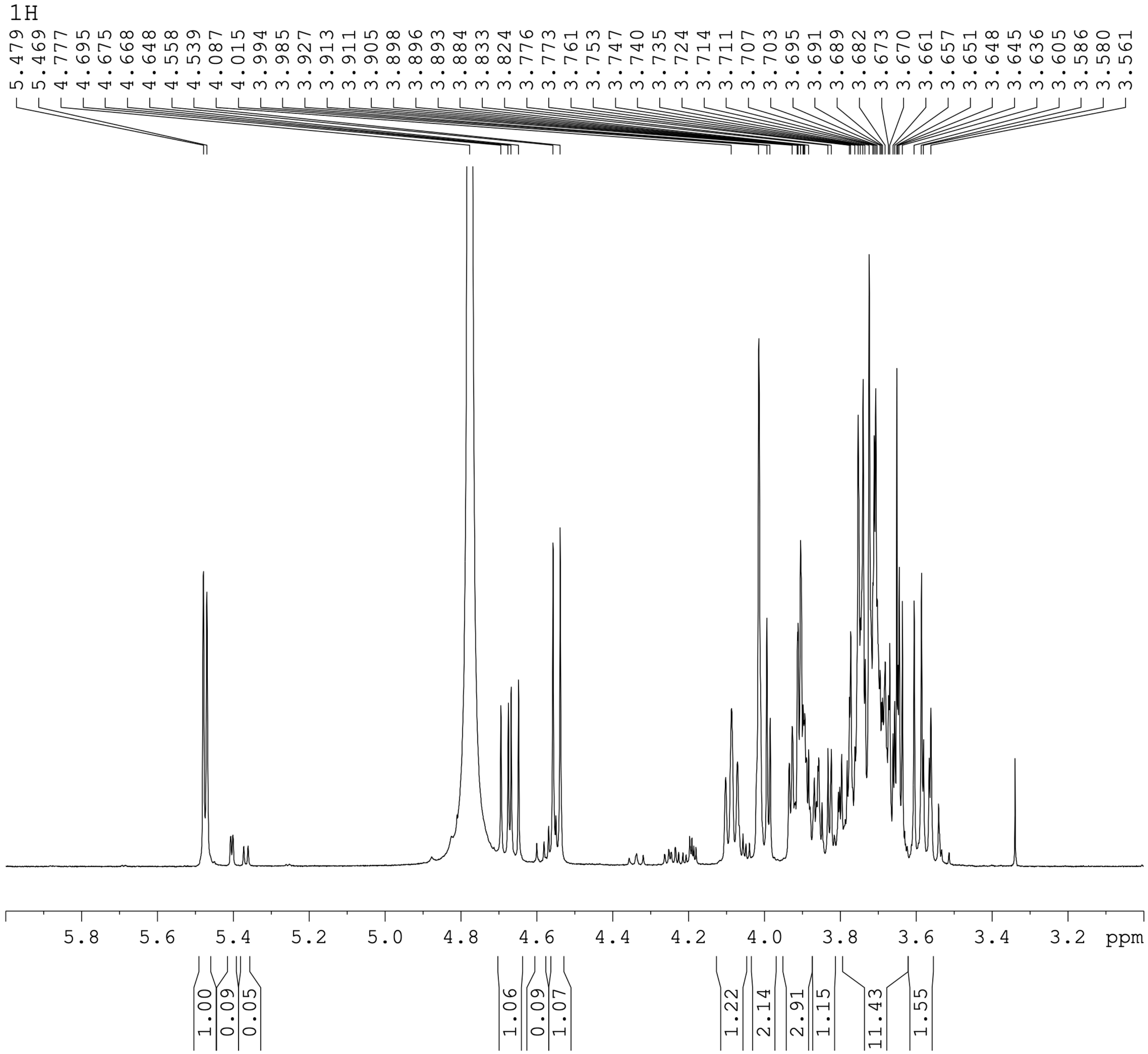
1H
5.479 5.469 4.777 4.695 4.675 4.668 4.648 4.558 4.539 4.087 4.015 3.994 3.985 3.927 3.913 3.911 3.905 3.898 3.896 3.893 3.884 3.833 3.824 3.776 3.773 3.761 3.753 3.747 3.740 3.735 3.724 3.714 3.711 3.707 3.703 3.695 3.691 3.689 3.682 3.673 3.670 3.661 3.657 3.651 3.648 3.645 3.636 3.605 3.586 3.580 3.561
5.8 5.6 5.4 5.2 5.0 4.8 4.6 4.4 4.2 4.0 3.8 3.6 3.4 3.2 ppm
1.00 0.09 0.05 1.06 0.09 1.07 1.22 2.14 2.91 1.15 11.43 1.55

13C

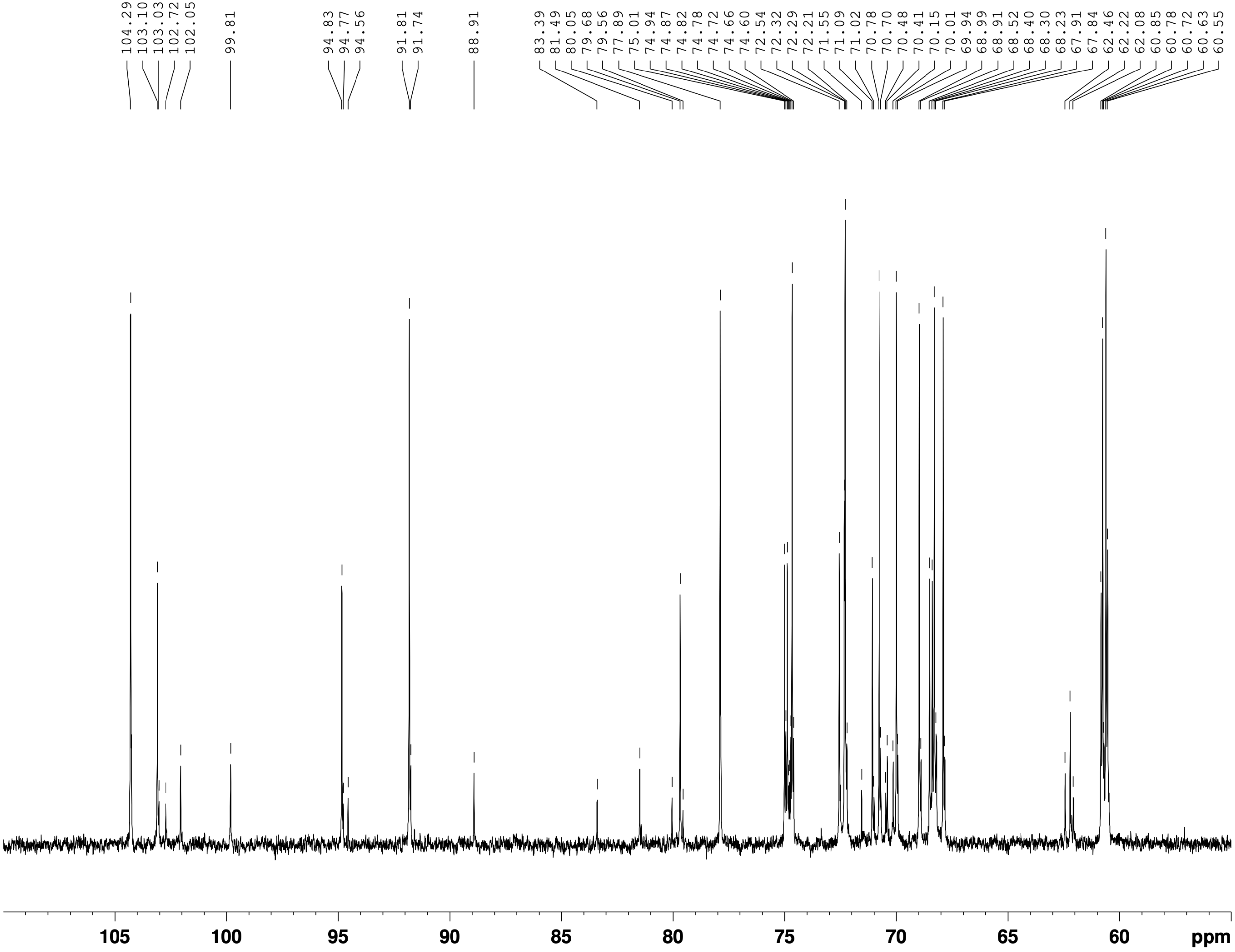
104.29
103.10
103.03
102.72
102.05
99.81
94.83
94.77
94.56
91.81
91.74
88.91
83.39
81.49
80.05
79.68
79.56
77.89
75.01
74.94
74.87
74.82
74.78
74.72
74.66
74.60
72.54
72.32
72.29
72.21
71.55
71.09
71.02
70.78
70.70
70.48
70.41
70.15
70.01
69.94
68.99
68.91
68.52
68.40
68.30
68.23
67.91
67.84
62.46
62.22
62.08
60.85
60.78
60.72
60.63
60.55
105
100
95
90
85
80
75
70
65
60
ppm

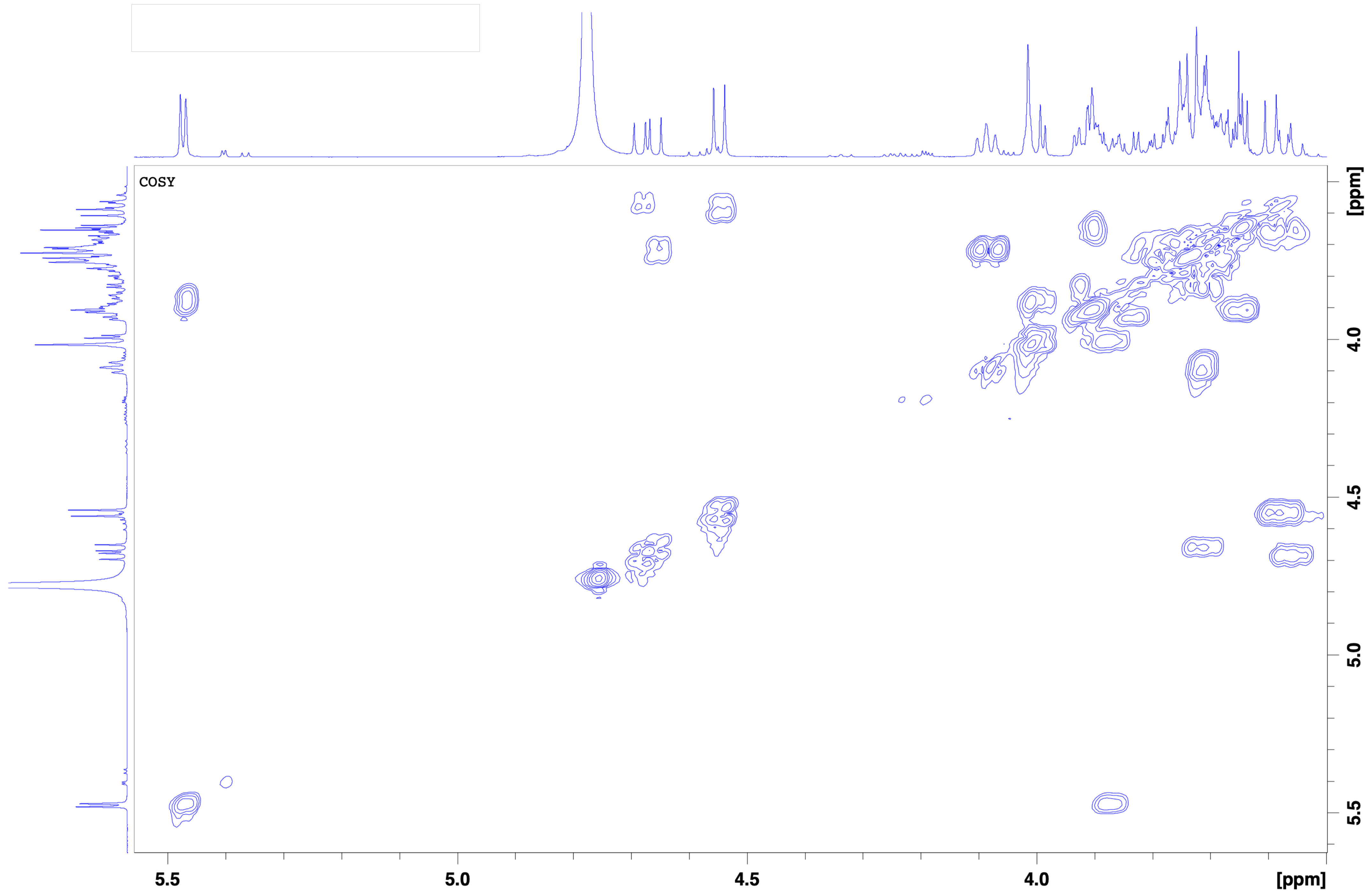
COSY
5.5
5.0
4.5
4.0
[ppm]
4.0
4.5
5.0
5.5
[ppm]

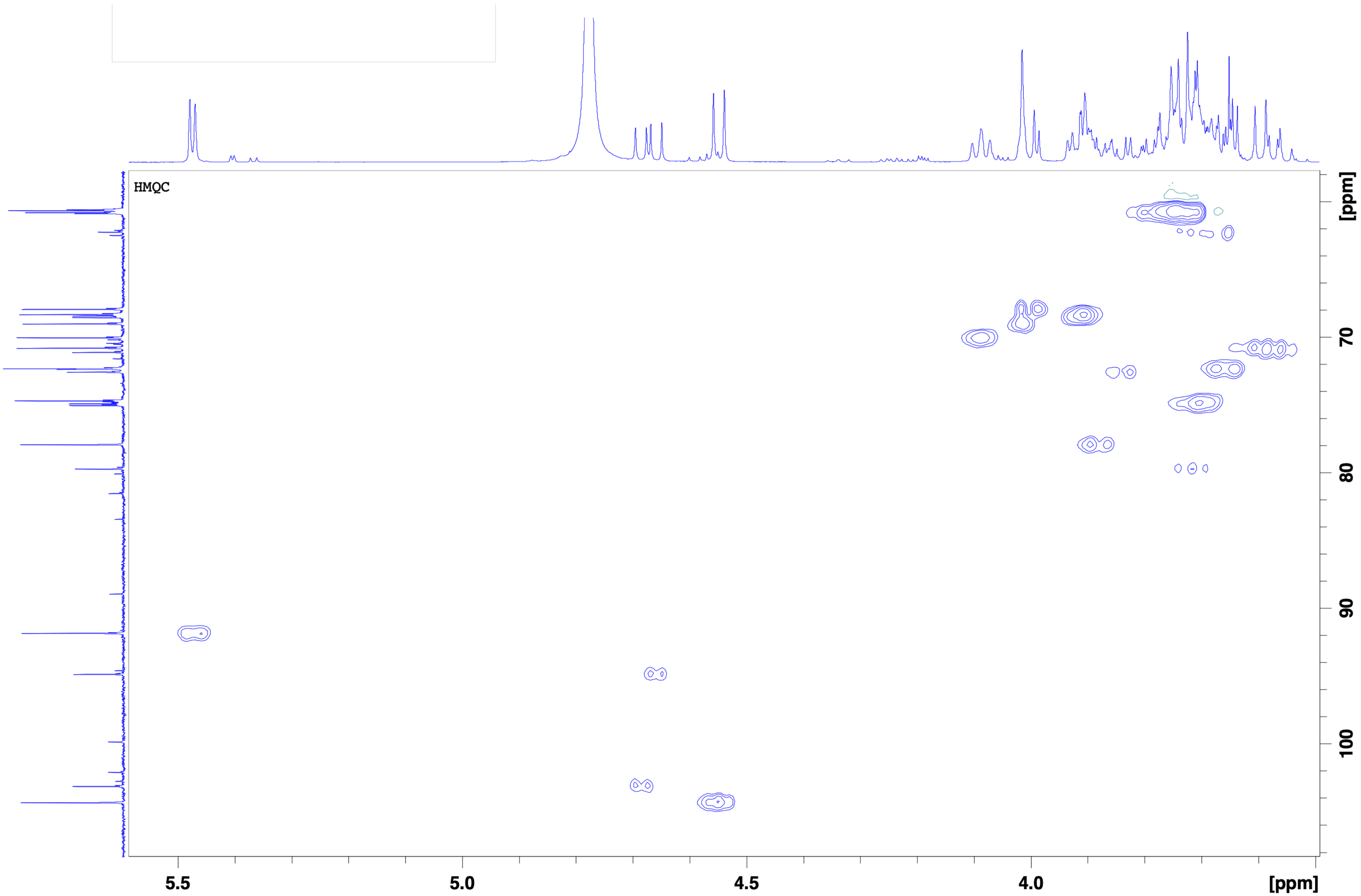

HMQC
5.5
5.0
4.5
4.0
[ppm]
70
80
90
100
[ppm]

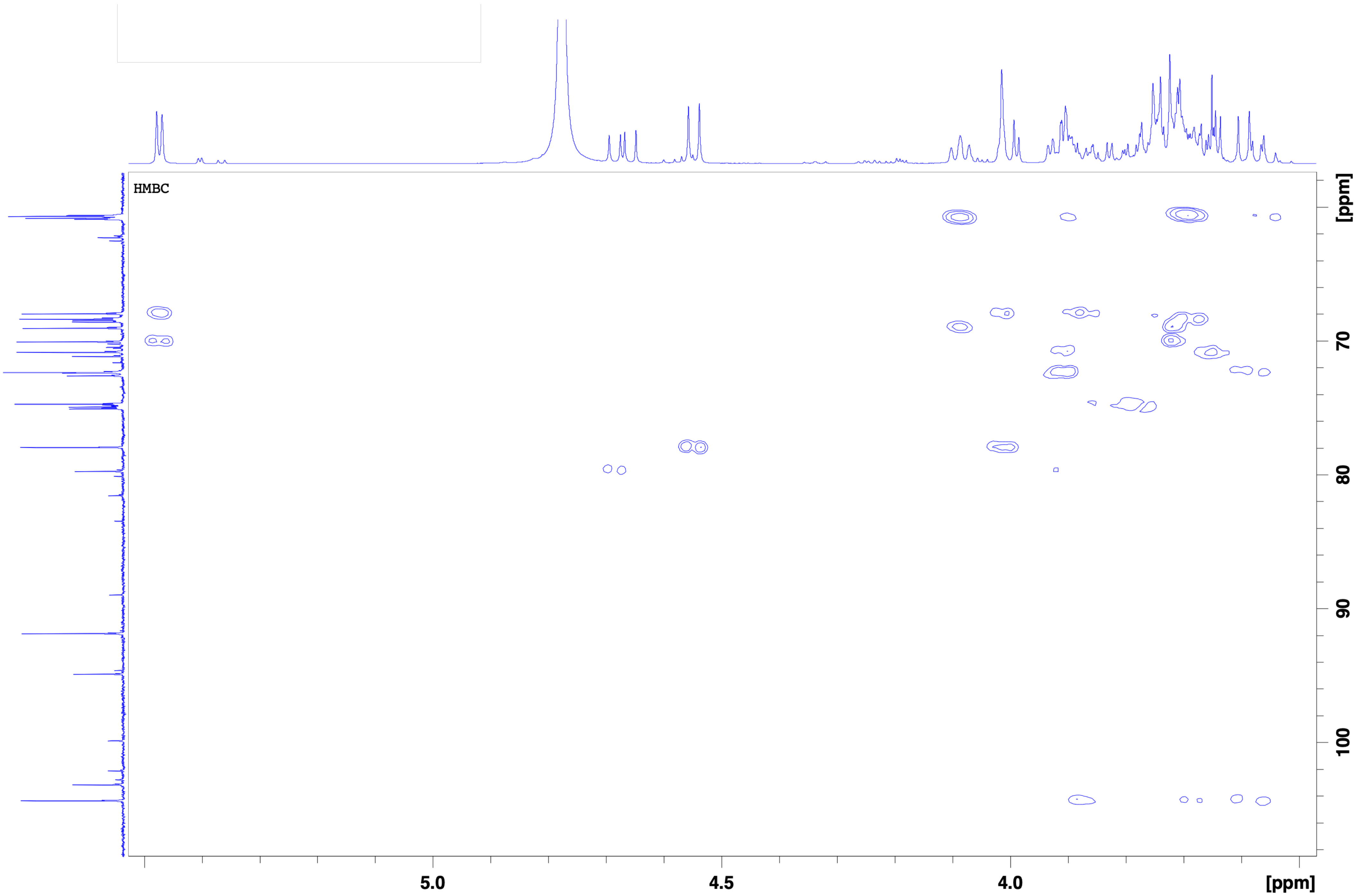
HMBC
[ppm]
70
80
90
100
5.0
4.5
4.0
[ppm]

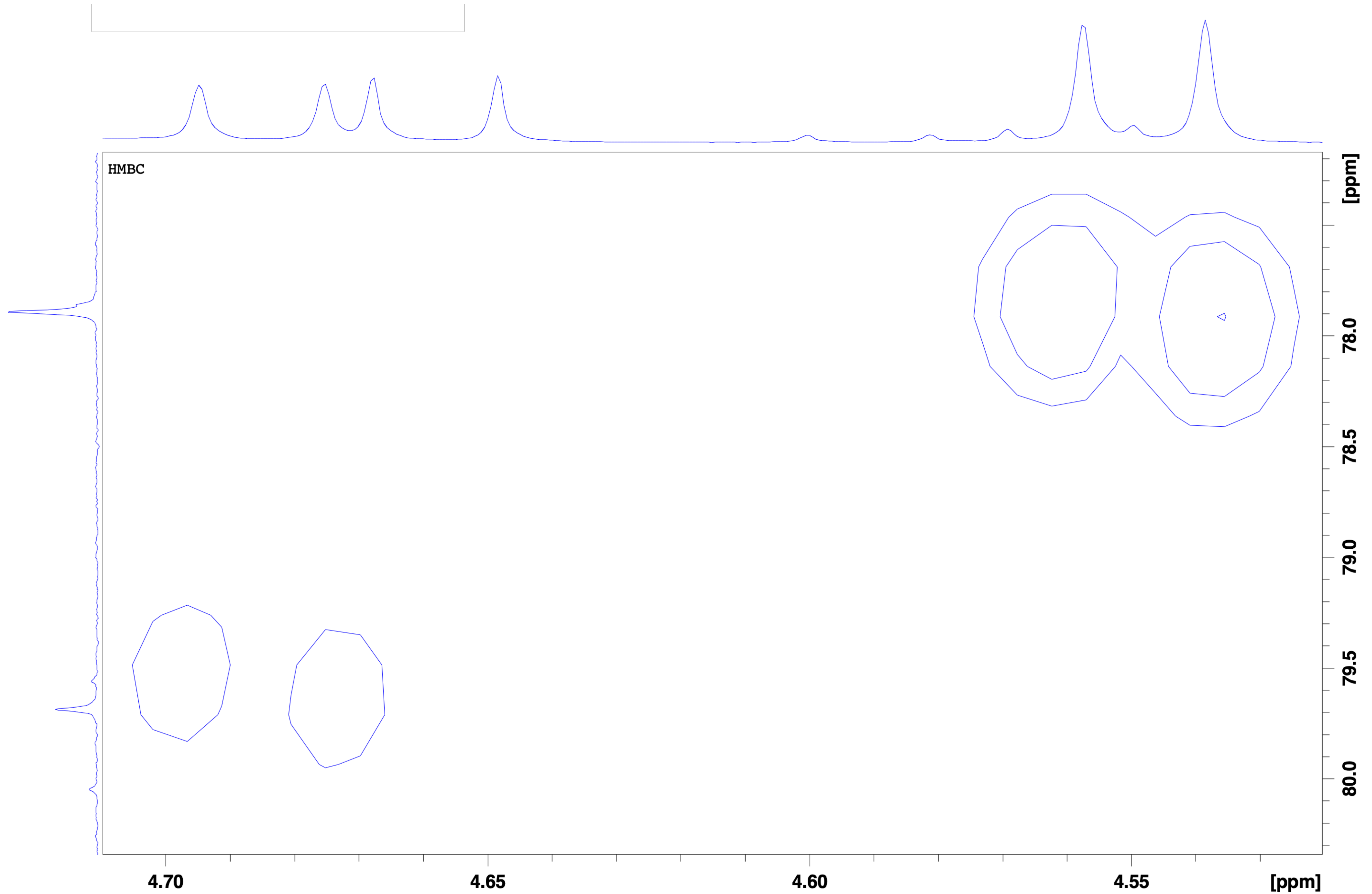
HMBC
[ppm]
78.0
78.5
79.0
79.5
80.0
4.70
4.65
4.60
4.55
[ppm]
